# High-resolution mapping of regulatory element interactions and genome architecture using ARC-C

**DOI:** 10.1101/467506

**Authors:** Ni Huang, Wei Qiang Seow, Julie Ahringer

**Affiliations:** The Gurdon Institute and Department of Genetics, Tennis Court Road, Cambridge CB2 1QN

**Author notes:** these authors contributed equally to this work (Ni Huang, Wei Qiang Seow).

## Abstract

Interactions between cis-regulatory elements such as promoters and enhancers are important for transcription but global identification of these interactions remains a major challenge. Leveraging the chromatin accessiblity of regulatory elements, we developed ARC-C (accessible region chromosome conformation capture), which profiles chromatin regulatory interactions genome-wide at high resolution. Applying ARC-C to *C. elegans*, we identify ~15,000 significant interactions at 500bp resolution. Regions bound by transcription factors and chromatin regulators such as cohesin and condensin II are enriched for interactions, and we use ARC-C to show that the BLMP-1 transcription factor mediates interactions between its targets. Investigating domain level architecture, we find that *C. elegans* chromatin domains defined by either active or repressive modifications form topologically associating domains (TADs) and that these domains interact to form A/B (active/inactive) compartment structure. ARC-C is a powerful new tool to interrogate genome architecture and regulatory interactions at high resolution.

## Results and Discussion

The development and application of chromosome conformation capture methods have been instrumental in shaping our understanding of genome topology^1, 2^. The basic premise is that if two regions of the genome are in close proximity within the nucleus, this information can be captured by the ability of such regions to be ligated together following DNA fragmentation. A diverse array of “C” methods have been developed, and their use has revealed interactions between regulatory elements, self-interacting topologically associated domains (TADs), and a “compartment” structure of the genome in which regions of similar activity interact like with like.

The Hi-C method uses high-throughput sequencing for genome-wide assay of chromatin interactions ^3^. In most Hi-C type methods, DNA is fragmented relatively uniformly using restriction or other enzymes to enable capture of interactions at all regions of the genome. This method maps large-scale genome architecture and domain structures, but it has insufficient resolution to widely identify interactions between specific regulatory elements. Such interactions have instead been mapped using oligo-based or other methods that capture targeted portions of a library, such as collections of promoters or regions bound by transcription factors^4^.

Chromatin at regulatory elements is known to be relatively accessible to nucleases, which has enabled their mapping using sensitivity to DNase I or Tn5 transposition (ATAC-seq)^5^). We reasoned that biasing fragmentation towards accessible chromatin would enrich a chromatin interaction library for interactions between regulatory elements, while still having sufficient information for interrogating larger scale architecture. This would avoid only obtaining regulatory information from user-defined regions and have the benefit of also mapping domain structure in the same experiment. Using this principle, we developed ARC-C (accessible region chromosome conformation capture).

The steps of ARC-C are illustrated in Figure 1a (see Methods). Nuclei fixed with formaldehyde are treated with a low concentration of DNase I (as in DNase hypersensitivity mapping) to enrich for cutting at accessible chromatin. Ends are then repaired and DNA ligated to join ends in close proximity. The library is then captured in nucleus using Tn5 tagmentation^6^, amplified, size-selected for inserts <400 bp, and paired-end sequenced. For data processing, we first identify “valid” high quality uniquely mapping read pairs. We then define “informative” read pairs that capture ligation events as those mapping >600 bp apart or to different chomosomes. Finally, we use “cis-informative” read pairs (those mapping to the same chromosome) to construct chromosome-wide contact maps and to call significant interactions following bias correction (Figure 1b-d; Supplementary Figure 1; see Methods).

**Figure 1.**
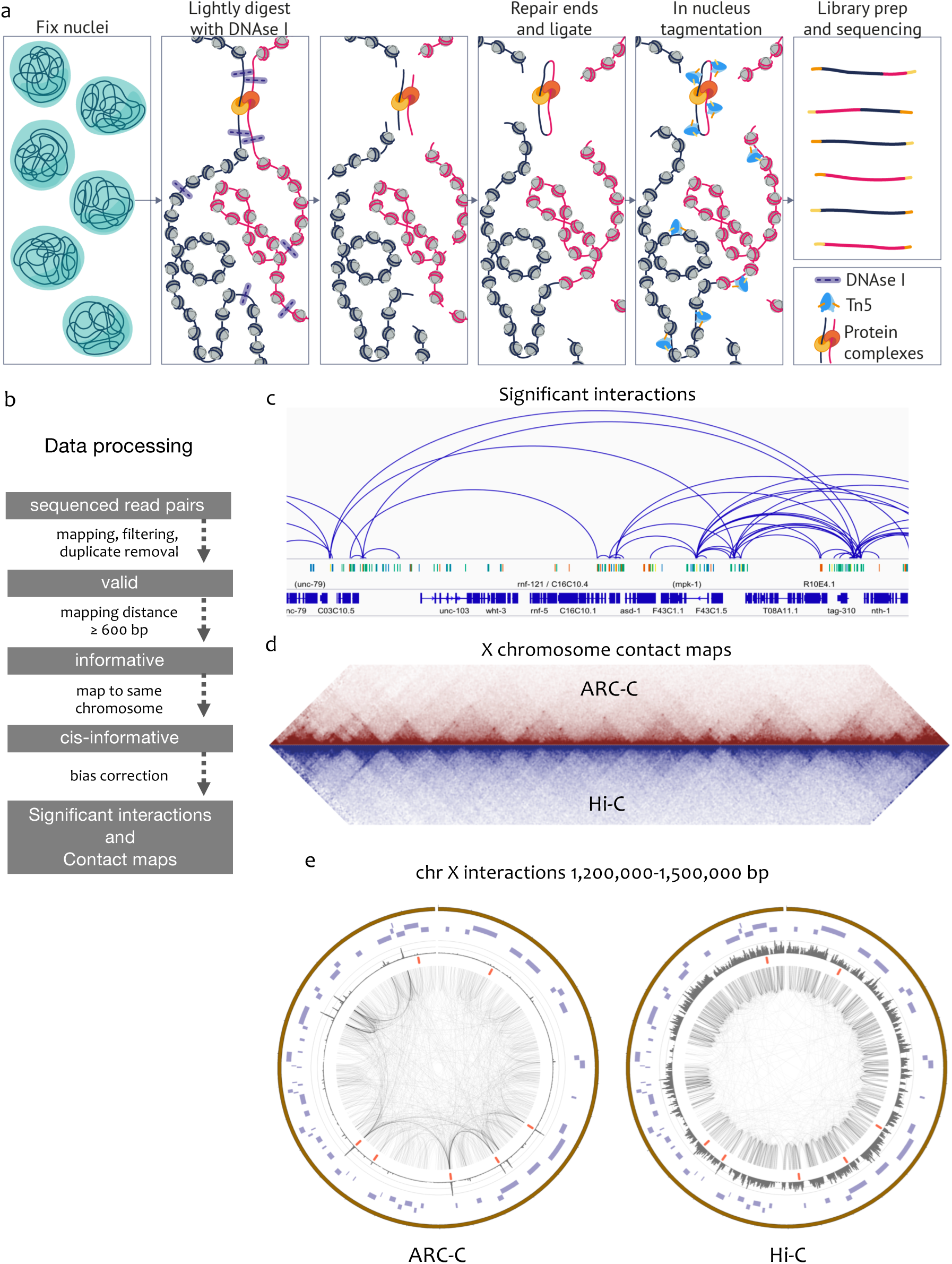
ARC-C method and comparison with Hi-C. (a) ARC-C cartoon. (b) Data processing steps. (c) IGV screen shot showing significant interactions in a 275kb window of chr III (4,075,000-4,300,822). Regulatory elements^23^(protein coding promoters, red; unassigned promoters, yellow; enhancers, green; unknown activity, blue) and genes are displayed below. (d) Comparison of ARC-C and Hi-C X chromosome contact maps. Hi-C data are from^7^. (e) Circos plots of cis-informative read pairs from ARC-C and Hi-C maps in 300kb region of chromosome X (1,200,000-1,500,000 bp). rex sites are indicated in red. Reads were downsampled in both plots for clarity.

We applied ARC-C to *C. elegans* L3 chromatin, preparing libraries from three biological and two technical replicates. Data from all replicates were highly concordant (Supplementary Figure 1). To increase the power to profile interactions, all cis-informative reads were pooled, resulting in 12 million reads pairs for analysis (Supplementary Figure 1). As expected, cis-informative read pairs were enriched at regulatory elements (43.7% of reads overlap regulatory elements, which comprise 21.1% of the genome).

Previous studies of *C. elegans* genome topology using Hi-C identified large self-interacting domains on the X chromosome regulated by the dosage compensation complex (DCC), which contain on average ~200 genes^7^. The DCC was shown to be enriched at recruitment element on X (rex) sites at domain boundaries, and 10 kb bins containing these sites were shown to physically interact with one another. We found that the ARC-C contact map recapitulated the X-chromosome domains mapped by Hi-C and more sensitively detected interactions between rex sites (Figure 1d, e). ARC-C contact maps from the autosomes are also highly similar to Hi-C maps (Supplementary Figure 2). These results show that ARC-C can profile domain large-scale domain architecture and suggest that it can improve detection of specific interactions between regulatory elements.

Previous Hi-C studies in *C. elegans* failed to identify smaller self-interacting topologically associating domains (TADs) similar to those seen in Drosophila and vertebrate genomes, which typically contain one to several genes^8-11^. Notably, the average *C. elegans* gene length of 5 kb (including intergenic regions) is similar to the resolution of Hi-C maps, which may have limited the ability to identify TADs.

We considered that genomic regions defined by chromatin modification domains might form TADs because in other animals, histone modification patterns within individual TADs is often relatively uniform^11^. In *C. elegans*, histone modification domains segment most of the genome into “active” domains of broadly active genes marked by H3K36me3 and other modifications associated with gene activity which alternate with domains of H3K27me3 covering genes that are inactive or have regulated expression^12, 13^. Both domain types have a median gene number of three, with median lengths of 14 kb for active domains and 19 kb for H3K27me3 domains^12^. To investigate whether these chromatin domains defined by histone modifications form TADs, we aggregated ARC-C signal across each type of aligned domain and their neighboring regions.

For both active and H3K27me3 domains, we found that ARC-C interaction frequency is higher within domains compared to interactions with neighboring chromatin, giving rise to a central square of higher signal. This indicates that the domains are spatially separated (Figure 2). We further observed that active and H3K27me3 domains have “compartment” structure, in which domains of the same type (active or H3K27me3) interact more frequently with each other than with domains of the opposite type (Figure 2). Supporting these analyses, we observed similar results using Hi-C data (Supplementary Figure 4). We conclude that *C. elegans* active and H3K27me3 chromatin domains form TADs and that these TADs are organized into a compartment structure similar to A/B compartments of other animals.

**Figure 2.**
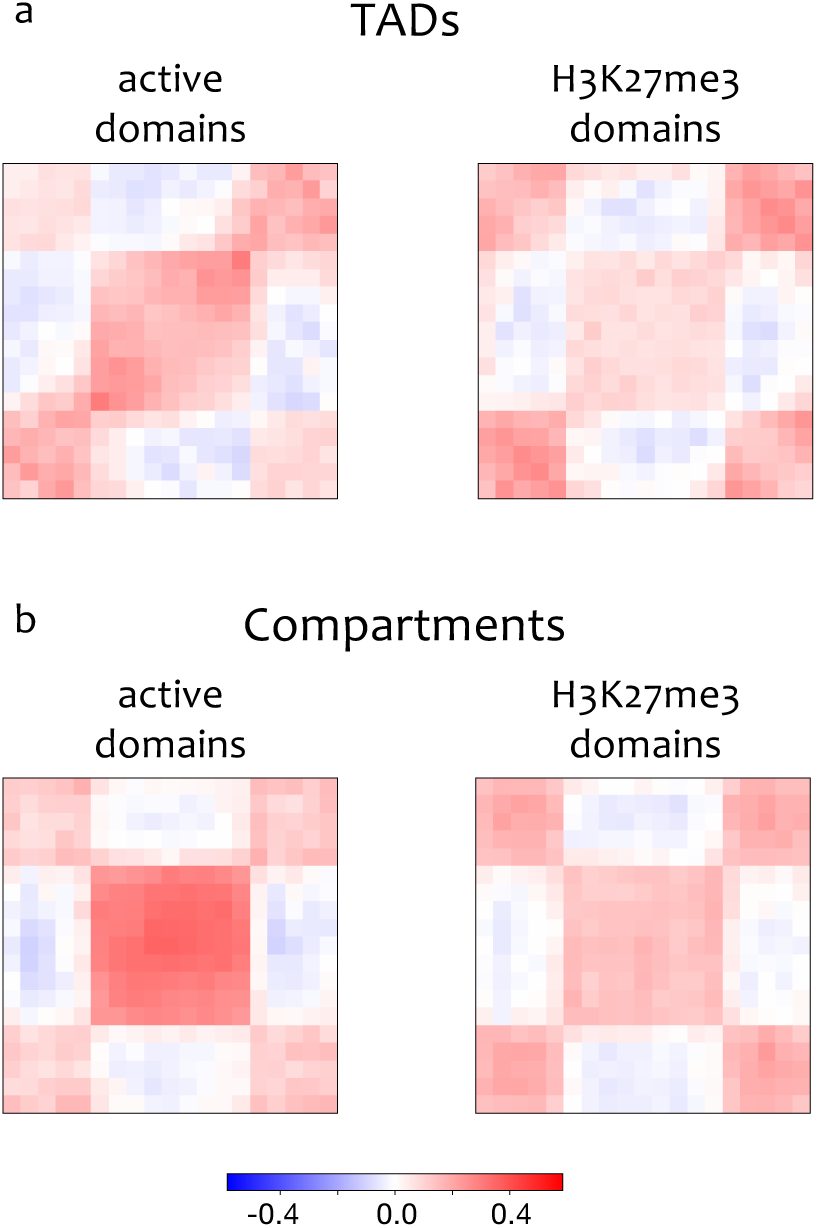
Chromatin modification domains form TADs that have compartment structure. “Active” and H3K27me3 (aka “regulated”) chromatin modification domains defined in Evans et al, 2016 were aligned and contact map signal aggregated in the aligned regions and neighboring regions. Higher signal in the central square shows enrichment for within domain interactions, indicative of TAD structure. Enrichment is 1.12 fold for active domains and 1.06 fold for H3K27me3 domains. (b) all possible pairs of inter-domain contacts in the range of 50kb to 2M were aligned and signal aggregated as in (a). Higher signal in the central square shows that domains interact more frequently with domains of the same type than with domains of opposite types, indicative of compartment structure. Enrichment is 1.23 fold for active domains and 1.1 fold for H3K27me3 domains. See Methods for details.

We next tested the ability of ARC-C to profile interactions between specific regulatory elements. Individual promoter and enhancer elements are small regions of DNA bound by transcription factors and from which transcription is initiated ^14^. To identify chromatin interactions between such elements at high resolution, we separated the genome into 500bp bins and identified those with significantly enriched interactions, taking into account distance and coverage biases (see Methods). This identified 14,992 chromatin interactions within a distance 1kb – 1Mb (Table S1). These involve 9733 regions, 95% of which overlap an annotated promoter or enhancer (Figure 3a; Table S2).

**Figure 3.**
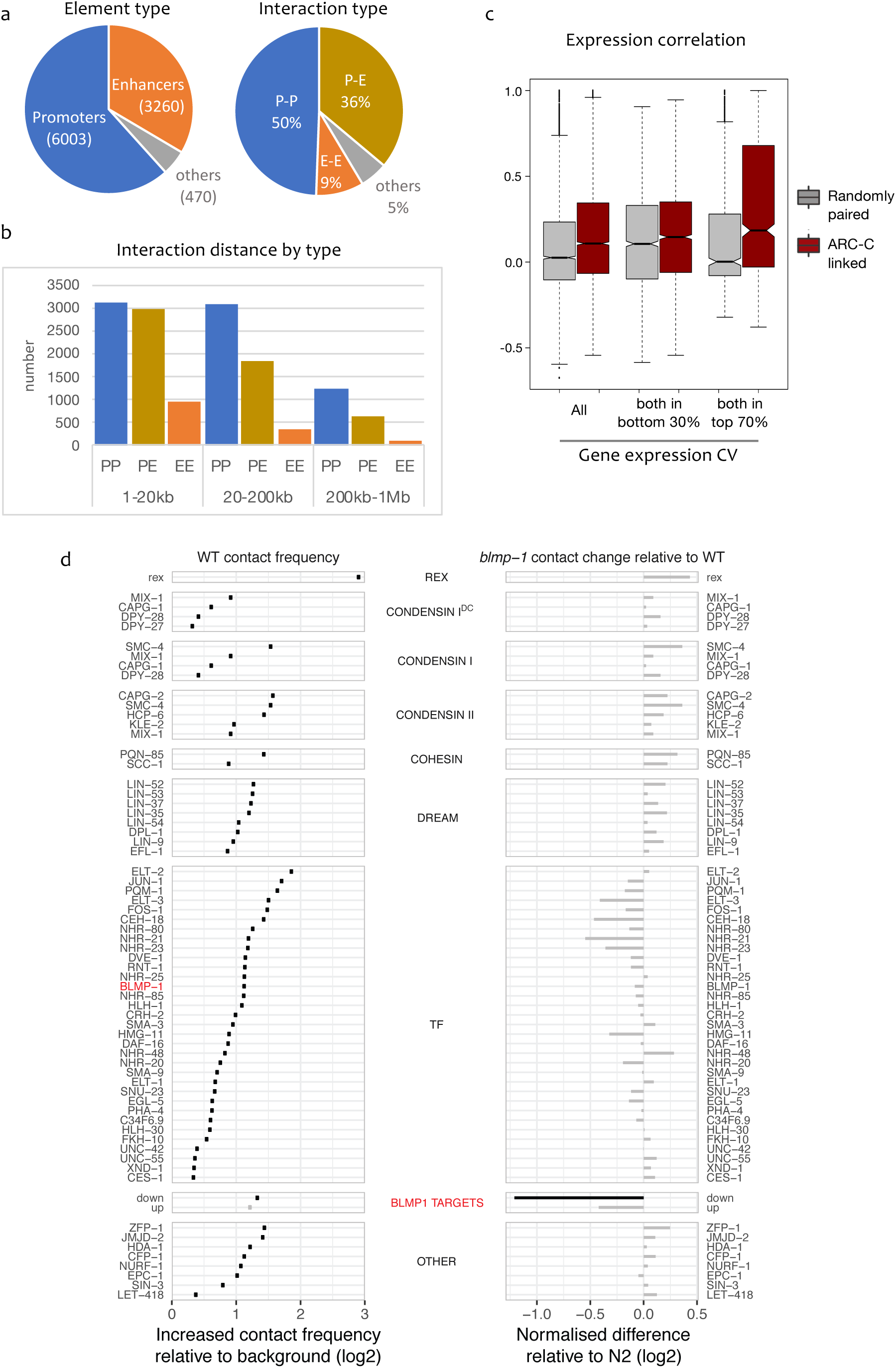
ARC-C defines significant interactions between regulatory elements, candidate regulators, and a role for BLMP-1 in mediating interactions. (a) Types of elements and types of interactions involved in the 14,901 significant interactions (b) Interaction distances of the significant interactions, separated by element type. (c) Expression correlation between pairs of genes with linked promoters compared to randomly paired genes in the set. Left, all pairs of linked genes (n=5142); middle, pairs for which both genes are in the bottom 30% of CV values (wide expression; n=2224); right, pairs for which both genes are in the top 70% of CV values (regulated expression; n=879). (d) Left, contact frequency between binding sites of indicated proteins. Right, change in contact frequency in *blmp*-*1* mutants compared to wild-type (log2). Interactions among *blmp*-*1* down targets are significantly reduced. See Methods for details and Table S5 for data.

Promoters are most prevalent among the significant interactions, accounting for 62% of interacting elements, and they are involved in 86% of the interactions (Figure 3a). Half of the significant interactions are relatively short range (within 20 kb), and in this size range there are similar numbers of P-P and P-E interactions (Figure 3b). However, at longer distances, promoter-promoter interactions predominate, which may indicate functional differences. We also observed that genes connected by promoter-promoter interactions had correlated gene expression (Figure 3c). The correlation is strongest for pairs with highly regulated expression (i.e, those with high coefficients of variation of gene expression (CV), suggesting that such genes are in proximity with each other when expressed.

To identify proteins that are candidates for mediating interactions in *C. elegans*, we screened for transcription factors and chromatin regulators for which binding sites show significantly enriched interactions in the ARC-C contact map (see Methods). Of 105 proteins tested, 60 chromatin regulators or TFs had this property (Figure 3d; Supplementary Figure 3; Table S3). Although the binding sites for only three proteins showed significant enrichment for interactions in the Hi-C map, the trends were similar to those observed using ARC-C (Supplementary Figure 3; Table S4). As expected from their role in mediating X chromosome interactions between rex sites^7^, components of the condensin I DC complex are in this set (Figure 3d). Other proteins of note are cohesin SCC-1 and loading factor PQM-85/NIPBL, consistent with similar enrichment in mammals and with the role of cohesin in loop formation^15, 16^. We also found enrichment for subunits of the condensin II and Retinoblastoma/DREAM complexes, 33 transcription factors, and other chromatin regulators (Figure 3d). These proteins are strong candidates for involvement in mediating chromatin interactions.

To evaluate the ability of ARC-C to detect changes in regulatory interactions, we chose to analyse BLMP-1, one TF for which binding sites significantly interact (Figure 3d). *blmp*-*1* encodes a spatially restricted transcription factor important for hypodermal, vulval, and gonadal development^17-19^. We performed ARC-C in L3 stage *blmp*-*1* mutants and compared chromatin interactions at pairs of TF and chromatin regulator binding sites with those of L3 stage wild-type. For this analysis, we also included direct BLMP-1 targets, defined as the subset of BLMP-1 binding regions associated with a gene that was upregulated or downregulated in *blmp*-*1* mutants. In wild-type, interaction strength among the up or down BLMP-1 target regions is similar to that of the full set of BLMP-1 binding sites (Figure 3d; Table S5). Strikingly, the only sites to show significantly reduced interactions in the ARC-C map from *blmp*-*1* mutants were the BLMP-1 down regulated targets (Figure 3d). These results indicate that BLMP-1 mediates interactions between targets that require it for expression. Overall, the results show that ARC-C has power to map interactions between regulatory elements at 500bp resolution and to detect changes in mutants.

In its present form, ARC-C works well for profiling chromatin interactions in relatively small genomes, as sequencing 200 million fragments per duplicate library produces enough cis-informative read pairs for profiling architecture and regulatory element interactions. However, for application to larger mammalian genomes, an enrichment step for ligation events (e.g., through biotin tagging) would be necessary.

Here we used ARC-C in whole animals, so the cell types from which the detected interactions came are unknown. In addition, interactions that occur in a small number of cells are likely to have been missed. The future application of ARC-C to specific purified cells would address these issues, allowing *in vivo* investigation of cell-type specific architecture.

In conclusion, ARC-C provides a new ability to assay genome topology together with high-resolution mapping of regulatory interactions in a genome-wide assay, which should accelerate studies of transcriptional regulation and the relationship with genome architecture.

## Methods

### Worm strains and culture

*C. elegans* strains were maintained at 20C as previously described^20^. The following strains were used: Bristol N2 (wild-type), YJ55 *blmp*-*1(tm548).*

### Worm growth

Strains were grown in liquid culture at 20°C using standard S-basal medium with HB101 bacteria. Animals were first grown to the adult stage, bleached to obtain embryos, and the embryos hatched without food in M9 buffer for 24 hrs at 20°C to obtain synchronized starved L1 larvae. L1 larvae were grown in a further liquid culture at 20°C then harvested at the L3 stage. Worms were collected, washed in M9 buffer, floated on sucrose, washed again in M9, then frozen into small pellets by dripping worm slurry into liquid nitrogen, which was stored at −80°C until use.

### ChIP-seq

JMJD-2, SCC-1, ZFP-1, BLMP-1, TOP-2, and SIN-3 chromatin immunoprecipitations in L3 larvae and library preparations were conducted as in^21^. Antibodies used were JMJD-1 (Q3951; Novus), SCC-1 (Q0835; Novus); ZFP-1 (Q2059; Novus), BLMP-1(Q2919, this study), TOP-2 (Q5515, this study), and SIN-3 (Q6013; ^22^. Peaks were called for each replicate separately as in ^23^ except for BLMP-1, which used MACS2^24^. Replicate peaks were combined using IDR^25^.

### Differential expression analysis of *blmp*-*1* mutant

Raw RNA-seq data of wild-type and *blmp*-*1* mutant at L3 stage were obtained from GEO (GSE55225)^17^. Reads were aligned to genome assembly ce10 with gene annotation WS235 using STAR^26^. Read counts per gene were generated by featureCounts and genes differentially expressed in *blmp*-*1* relative to wild-type identified using DESeq2^27^ requiring adjusted p < 0.05 and absolute fold change > 1.5. Differentially expressed genes for which a BLMP-1 ChIP-seq peak overlapped one of its assigned regulatory elements were defined as upregulated “up” or downregulated “down” BLMP-1 targets.

### ARC-C library preparation

Frozen worm pellets were ground into a fine powder in which worms were broken into ~10 fragments. 1 ml of worm powder was then fixed in 10 ml of 1% formaldehyde in PBS at room temperature (RT) for 10 min with gentle shaking then quenched for 5 min with a final concentration of 125mM glycine. Fixed worm fragments were then washed with Buffer A (340mM sucrose, 15mM Tris-HCl pH7.5, 2mM MgCl2, 0.5mM spermidine, 0.15 mM spermine, 1mM DTT, protease inhibitors), resuspended in 7 ml Buffer A, then the material dounced 20 strokes in a 7 ml stainless steel tissue grinder (VWR-#432-5005). The material was spun at 100g for 5 minutes and the supernatant containing nuclei was transferred to a new tube and kept. The pellet was resuspended in Buffer A and again dounced with 20 strokes. After spinning, the two supernatants containing nuclei were pooled.

Three aliquots of 10 million fixed nuclei were spun down at 1,000g and resuspended in 200ul of 1x DNase buffer (Roche) and chromatin was digested with 25, 50, or 100U/ml DNase I at 25C for 10min. The reactions were then quenched with a final concentration of 25mM EDTA and 5mM Tris pH 7.5. Nuclei were washed twice with 1ml of ice-cold Nuclear Washing Buffer (340mM sucrose, 15mM Tris-HCl pH7.5, 25mM EDTA, 0.5mM spermidine, 0.15mM spermine, 1mM DTT, protease inhibitors). Nuclei were then resuspended in 100ul of end repair master-mix (10ul 10x NEB End Repair Buffer, 5ul NEB End Repair Enzyme Mix, 85ul H2O) and incubated at 20C for 30min with rotation. Thereafter, 400ul of ligation master-mix was added (40ul 10x ligation buffer, 5ul T4 DNA ligase [400,000 units/ml], 355ul H2O) and the mixture was incubated at 4C overnight with rotation. Nuclei were pelleted, resuspended in 50ul of tagmentation master-mix (22.5ul H2O, 25ul 2x Nextera TD buffer, 2.5ul Nextera Tn5 transposase) and incubated at 37C for 30 min. 5ul of 1% SDS and 2ul of NEB Proteinase K [800U/ml] was added and left at 65C for 15 min before DNA from the mixture was purified using Qiagen MinElute columns. Large fragments (>500bp) were removed from the purified DNA using two rounds of 0.6vol AMPure XP beads. Prior to PCR amplification, a test qPCR amplification using 1/20^th^ of the input DNA was performed and the cycle number for PCR was determined by the midpoint of the exponential phase of the amplification curve. The resultant DNA was amplified with NEBNext Ultra II Q5 Master-mix under the following PCR conditions: 72 °C for 5 min, 98 °C for 30 s, and cycling at 98°C for 10s, 63°C for 30s and 72°C for 1min using the determined cycle number. The library was size-selected with AMPure XP beads to a final range of 200-700bp (insert size: ~70-570bp). Libraries made from nuclei digested with different concentrations of DNase I were quality tested by qPCR at the gap-3 promoter. DNAse I over- or under-digestion led to poor enrichment of regulatory elements. Good libraries generally had *gap*-*3* promoter enrichment values of >4-fold. Libraries were sequenced on the Illumina platform paired-end (100bp). Four libraries were sequenced from wild-type L3 larvae (three biological and two technical replicates) and two biological replicate libaries were sequenced for *blmp*-*1* L3 larvae.

### Processing ARC-C data

ARC-C libraries were sequenced 62 bp-150 bp (100 bp for most samples) from both ends. Adapter sequence was trimmed by cutadapt and sequences with under 20 bp remaining were removed. Each sequenced end was aligned independently to the ce10 reference genome using BWA mem^25^ which allows split-read-alignment using the default parameters. The two aligned ends (or the 5’ segment for split alignments) were then paired. We required both ends of a pair to align uniquely and with high confidence (mapping quality ≥ 30 and number of mismatches ≤ 2) to the nuclear genome and outside modENCODE backlisted-regions. PCR duplicates were next removed by sambamba markdup. The remaining read pairs were regarded as *valid* read pairs. Valid read pairs mapping to different chromosomes or greater than 600bp apart on the same chromosome were regarded as *trans*- or *cis*-*informative* read pairs, respectively. The 600bp threshold was established by comparing the proportions of the four possible end alignment orientation configurations (forward-forward, forward-reverse, reverse-forward and reverse-reverse) as a function of mapping distance. The vast majority of pairs mapping under 500 apart bp were in the forward-reverse configuration (non-ligated fragments), whereas above 600bp the proportions were stably at ~25% each.

Contact maps were made from informative read pairs by binning the genome into fixed-width non-overlapping bins and counting the number of read pairs between each pair of bins. The maps were then normalised by matrix balancing using the Knight-Ruiz algorithm^28^. For aggregated contact analysis (see below), the map was further divided by the spline-smoothed average contact frequency given distance from the diagonal to remove background slope of contact frequency.

### Processing Hi-C data

Raw FASTQ files of wild-type mixed embryo Hi-C data^7^ were downloaded from SRA (SRX77040) and processed using HiCUP v0.5.9^29^, which filters for same fragment (circularised, dangling ends, internal), religation, wrong size, contiguous sequence and removed duplicate read pairs. We additionally required mapQ>=30 and number of mismatches <= 2 from both reads, that none of the reads overlapped modENCODE blacklisted regions, and a minimum distance of 600bp between the two read pairs to be consistent with the processing of ARC-C data. In the end, 25,460,294 read pairs passed all filters, out of which 17,100,808 have both reads mapping to the same chromosome.

### Calling significant interactions

We first segmented the genome into bins of ~500bp. We first took annotated regulatory elements^23^ (n=42,245) and expanded them to 500bp or until neighbouring intervals began to touch; a small number of elements that were within 100bp were merged first. The rest of the genome was covered with evenly placed 500bp nonoverlapping fixed-width intervals, hence the entire genome was covered by a combined set of 192,257 intervals of average size (494 bp). We used this procedure instead of generating fixed non-overlapping bins to avoid individual REs being split into two bins.

Before assessing the significance of chromatin interactions in Hi-C or other chromatin interaction datasets, inherent bias resulting from uneven coverage and physical proximity need to be accounted for. A difference in ARC-C data compared to that of Hi-C is the specific over-representation of open chromatin. The observed enrichments at these regions are a compound effect of enriching the regions themselves and of their being enriched by virtue of linkage to another region of open chromatin (true signals). To address this and avoid normalising out interaction signals, we developed an approach similar to that used for Capture Hi-C, which has similar coverage biases and used “off-peak” interaction frequency to account for coverage^30^. We modelled the number of read pairs linking two intervals as a random variable following a binomial distribution parameterised by an expected contact frequency determined by unevenness of coverage and distance between the interval^31^:

For every interval *i*∈*I*, the number of cis-informative read pairs *c*_*cis*,*i*_ were counted. Intervals in the top 10% of the coverage distribution were regarded as peaks and intervals in the bottom 10% were removed. An off-peak cis-informative coverage *c*_*offpeak*,*i*_ was calculated for every kept interval, counting the number of contacts not involving peak intervals. We calculated a scaling factor for the interval’s representation/visibility as 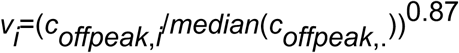 (see below for derivation of the exponent). Chromosome-wide average distance-dependent contact frequency *F*(*d*) in the distance range of 1 kb to 1 Mb was modeled by fitting a spline function in a two-pass process (following Fit-Hi-C^31^). For every pair of intervals with a distance between 1kb to 1Mb, an expected contact frequency was calculated given the distance and the visibility of each interval as 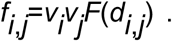 Given the total number of cis-informative contacts (*N*) of the chromosome, we considered a null distribution in the form of a binomial, where the observed number of contacts, 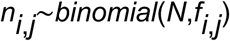. Significant interactions were called at an FDR level of 0.05 and were post-filtered requiring support by more than 5 read pairs.

The appropriate correction factors for adjusting the representation bias, *X*, should be able to transform the unnormalised contact matrix *A* into a normalised contact matrix *B* by *B*=*diag*(*X*)^∗^*A*^∗^*diag*(*X*), such that each row or column of *B* sums up to the same value, thus eliminating the unevenness of representation across different bins. It has been well-established that a correct set of factors can be found using the method “matrix-balancing” (MB)^15, 32^ and efficient algorithm has been developed^15, 28^. Our goal is to find correction factors that enable transformation into a normalised contact matrix in which each bin has the same off-peak coverage. As an efficient algorithm for solving this problem at high resolution is not yet available yet, we aimed to find an approximate solution. Two measures, namely the reciprocal of coverage (*c*) and square root coverage (*c*^0.5^) have been proposed for use in the place of MB-derived correction factors^15^. It was reported that the former can over-correct whereas the latter gives good approximation to MB-derived correction factors^15^. We examined correlation between MB-derived correction factors and the reciprocal of different exponents of coverage (Supplementary Figure 5), and found that the reciprocal of *c* indeed over-corrects, but the reciprocal of square root coverage under-corrects. The most accurate approximation is achieved at around an exponent of 0.87. Therefore, we used the reciprocal of (*c*_offpeak_)^0.87^ as the correction factor.

### Aggregated contact analyses

A contact is defined as the region in the contact map that connects a pair of genomic locations. Aggregated contact analysis is a method of visualising the average contact frequency of a group of many contacts together with local contact frequency^15^. We applied this method to both small genomic regions, such as nuclear factor ChIP-seq binding sites (NFBS), and to larger intervals, such as chromatin domains. We normalised contact maps using matrix balancing^32^ to account for coverage bias and removed distance dependent background. In NFBS analysis, we used contact maps of 1kb resolution. For each NF, up to 50,000 contacts were randomly sampled from all possible cis-contacts among its binding sites within a distance range of 20kb to 1Mb, and local maps of 21 by 21 bins centered at the contacts were extracted and aggregated. For the case of BLMP-1 regulated targets, all possible cis-contacts between BLMP-1 binding sites that involves a BLMP-1 regulated target were aggregated (i.e. at least one end of the contact is at a BLMP-1 regulated target). The log2 fold change of the central point over the mean of the rest of the points in the aggregated map was calculated to measure the relative increase in contact frequency over local background. To assess statistical significance while controlling for accessibility and the distance between the pair of NFBSs, 1000 sets of random contacts were generated, each containing the same number of contacts with matching accessibility and distance as the NFBS contacts. Each of the 1000 random sets was aggregated and a relative increase in contact frequency calculated in the same way as the NFBS set, forming a distribution of values against which the NFBS value was compared and a p-value generated, which was corrected for multiple testing using the FDR method. Datasets are from^21,33-38^. Processed and curated modENCODE ChIP-seq peaks were obtained from^23^. We removed intervals that are “HOT” (highly occupied target) as such binding events are thought to represent nonsequence specific TF binding or ChIP artifacts^36, 39^. This was defined as the top 20% of peak intervals ranked by the number factors in which the interval is called (effectively removing intervals called in 12 or more factors). We only considered datasets having at least 300 peaks following filtering.

For domain analyses, we used contact maps of 5kb resolution. The active and H3K27me3 domains are L3 domains from^12^. Regions annotated as active and border were merged to generate the active domains used here; the H3K27me3 domains are those termed “regulated.” We tested for TADs by assessing intra-domain contacts. For each type of domain (active or H3K27me3), maps of each contact region containing a domain of at least 5kb together with up to 25kb of the flanking domains of opposite type were extracted and aggregated. Where neighboring domains were < 25 kb, only the domain was extracted. The aggregated contact was scaled to a square of 9 by 9 bins and the flanking intervals were scaled to 5 bins wide. The log2 fold change of the mean of the central 9 by 9 square over the mean of the four neighbouring 5 by 10 rectangles on top, bottom, left and right was calculated to measure the relative increase in contact frequency with p-values generated by t-test. We tested for compartments using the same approach, by assessing all possible pairs of inter-domain contacts in the range of 50kb to 2Mb.

### Aggregated contact analysis of *blmp*-*1* mutant ARC-C data

We used the procedure described above for wild-type to measure the relative contact frequency between NFBSs in *blmp*-*1* ARC-C data. To normalize for a global linear reduction in open chromatin enrichment in the blmp-1 map, we we regressed out this effect by fitting *blmp*-*1* ACA measures as a linear function of the respective measures in wild-type (i.e. log2(blmp1_FC) = a + b ^∗^ log2(N2_FC). The residuals were used to measure the difference in contact frequency in *blmp*-*1* relative to wild-type. Statistical significance was assessed by 10000-time boot-strapping the distribution of residuals and p-values were adjusted by FDR (Table S5).

### Expression correlation between interacting promoters

For genes linked by promoter-promoter interactions, we calculated Pearson correlation coefficients of gene expression across cell types using data from^40^. For pairs involving bi-directional promoters, the gene pair with highest correlation was chosen. For each gene, we also calculated a coefficient of variation of gene expression (CV) across the cell types^40^ as a measure of tissue-biased expression.

Genes with similar expression across cell types have low gene expression CV values and those with tissue-biased expression have high CV values. In Figure 3c, we assessed expression correlation between all linked pairs of genes (n=5124), linked genes for which both were in the bottom 30% of all CV values (n=2224), and linked genes for which both were in the top 70% of CV values (n=879). To assess the statistical significance of the correlations, we generated control sets by sampling same number of random pairs using genes from the respective linked gene sets while requiring the distribution of distance between the random pairs to match that of the observed set. Difference in correlation between the observed and the random control set was tested using a t-test: all (p=7.77e-27), bottom 30% (p=4.38e-06), top 70% (p=5.69e-20).

## Acknowledgements

We thank K. Harnish for sequencing and R. Durbin for comments on the manuscript. The work was supported by Wellcome Trust Senior Research Fellowships to JA (054523 and 101863) and an A-star fellowship to WQS. We also acknowledge core support from the Wellcome Trust (092096) and Cancer Research UK (C6946/A14492).

## Figure legends

**Supplementary Figure 1.**
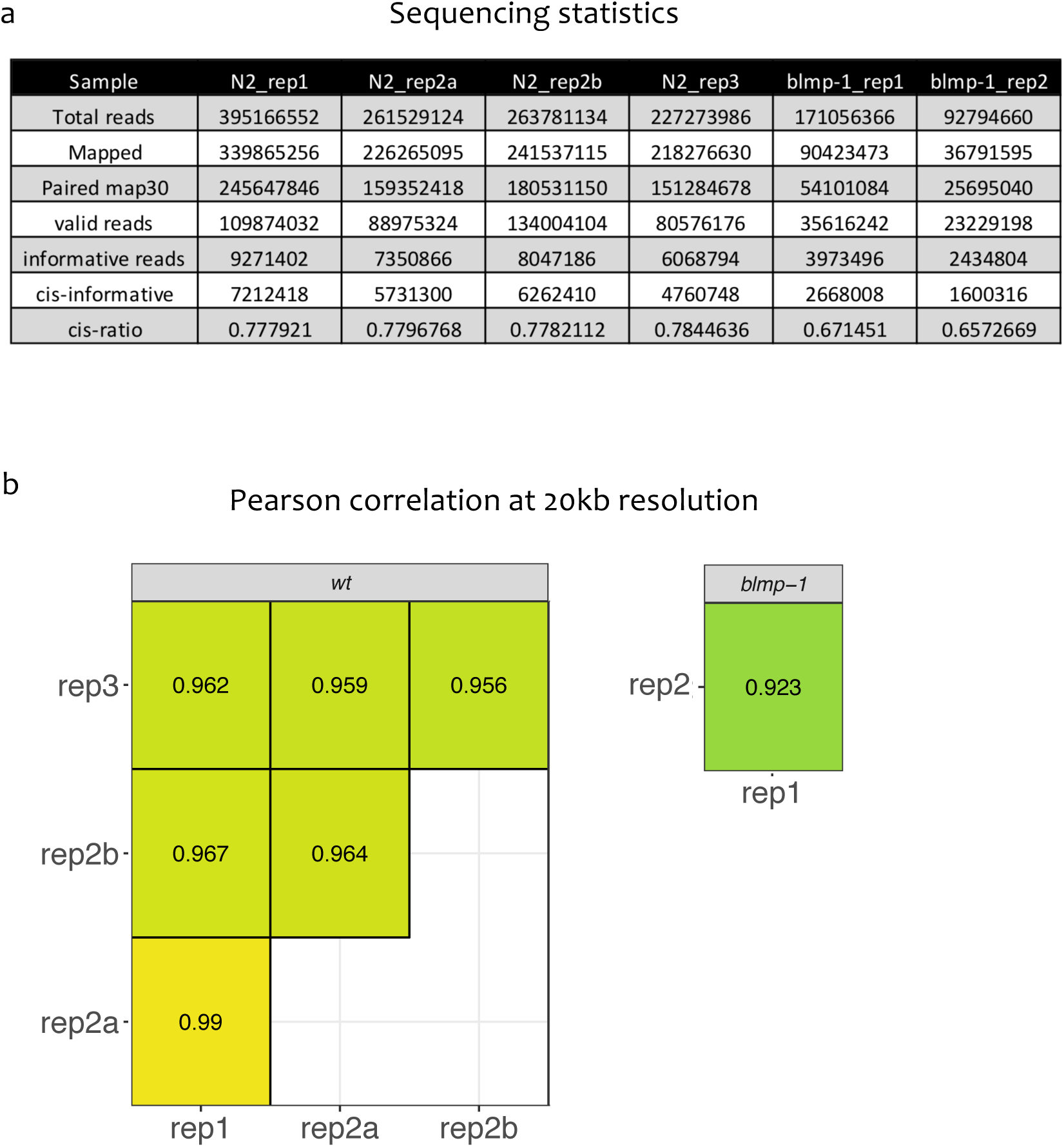
Sequencing statistics and replicate correlations. (a) Sequencing statistics of reads for each replicate. (b) Pearson correlation coefficients between replicates using 20 kb binned data.

**Supplementary Figure 2.**
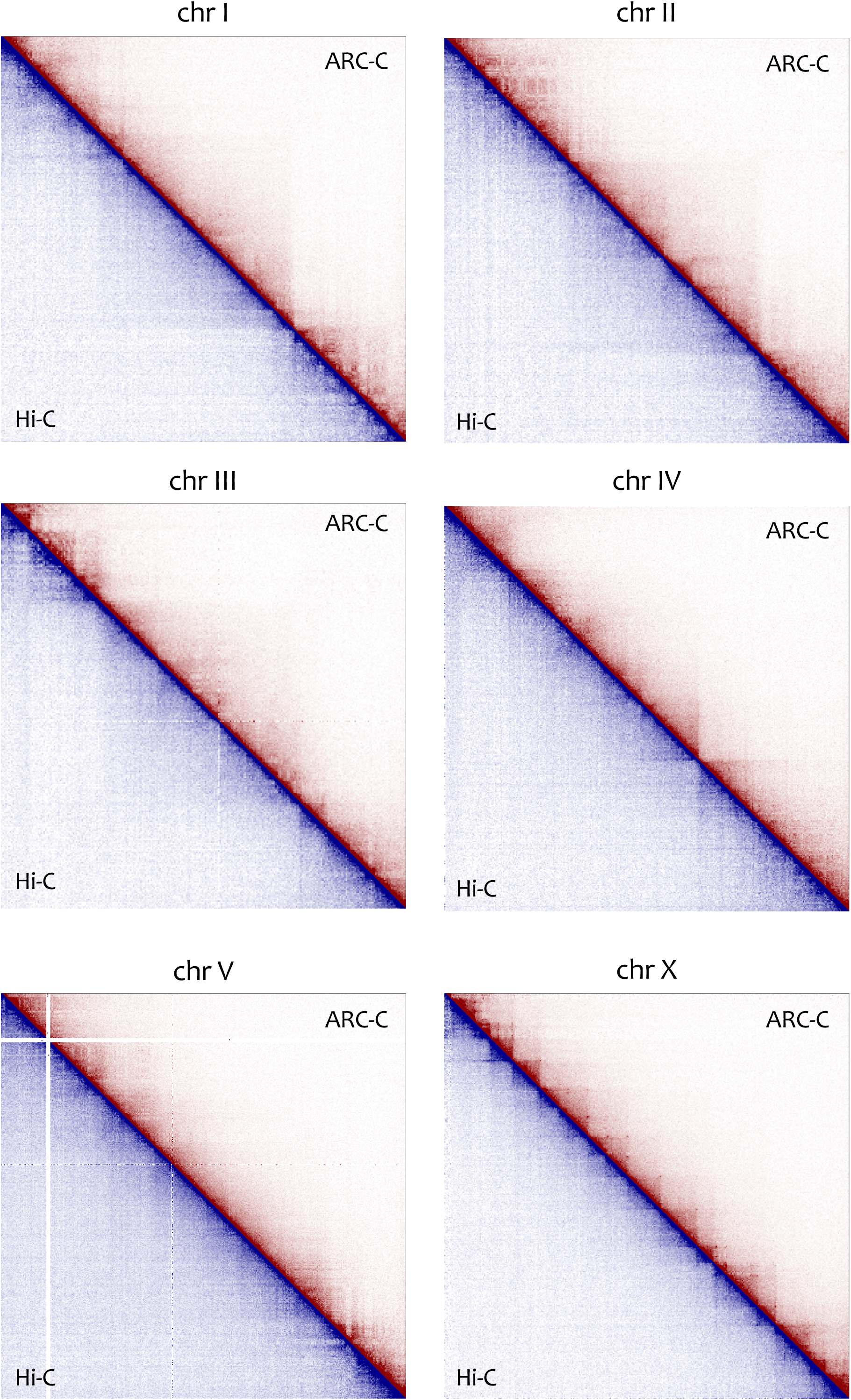
Comparison of ARC-C and Hi-C contact maps for each of the six *C. elegans* chromosomes at 50 kb resolution. Hi-C data are from^7^.

**Supplementary Figure 3.**
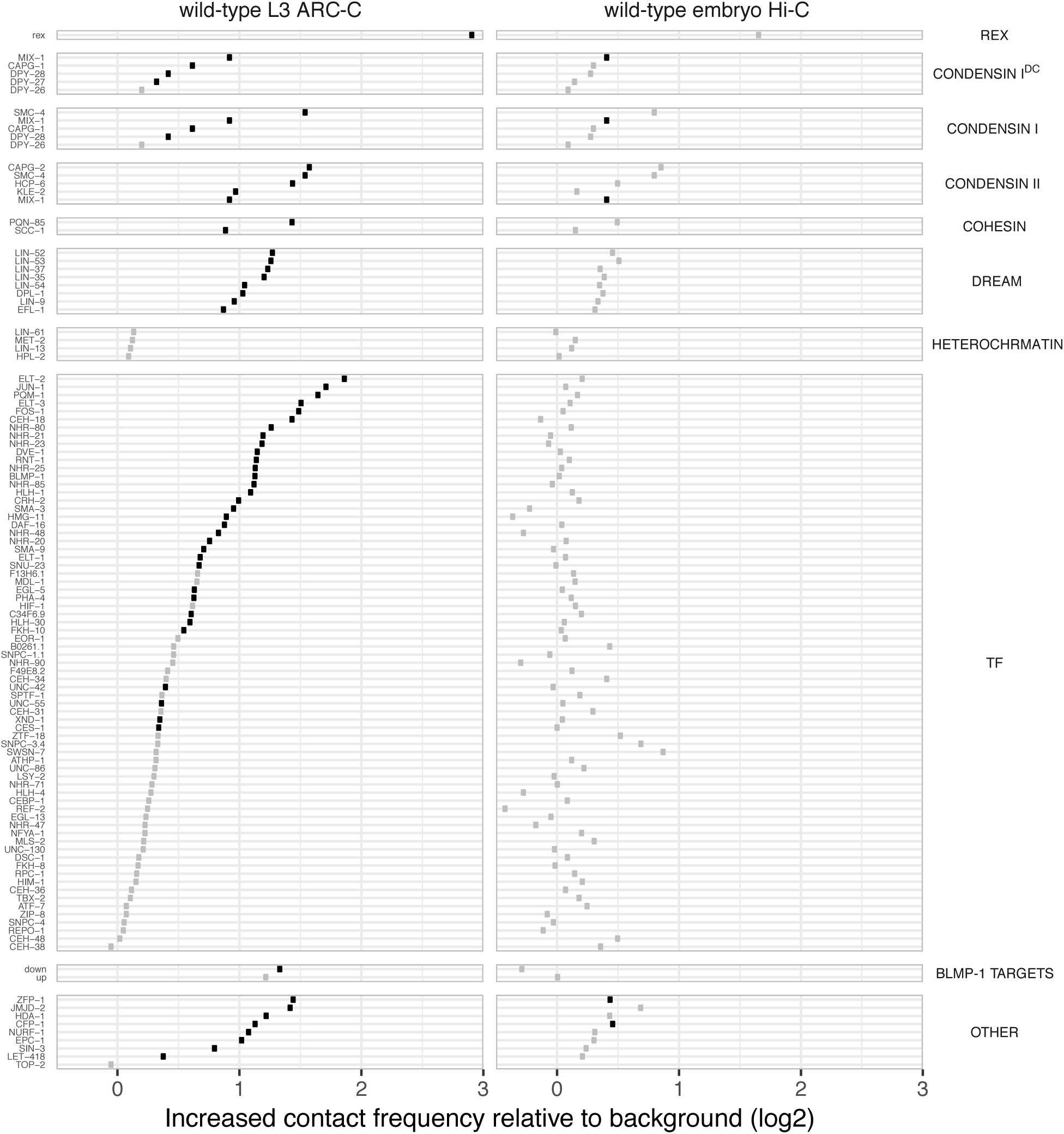
Comparison of contact frequency between binding sites of indicated proteins in ARC-C and Hi-C maps. Black circles indicate significant enrichment. Hi-C data are from Crane et al, 2014. See Methods for details.

**Supplementary Figure 4.**
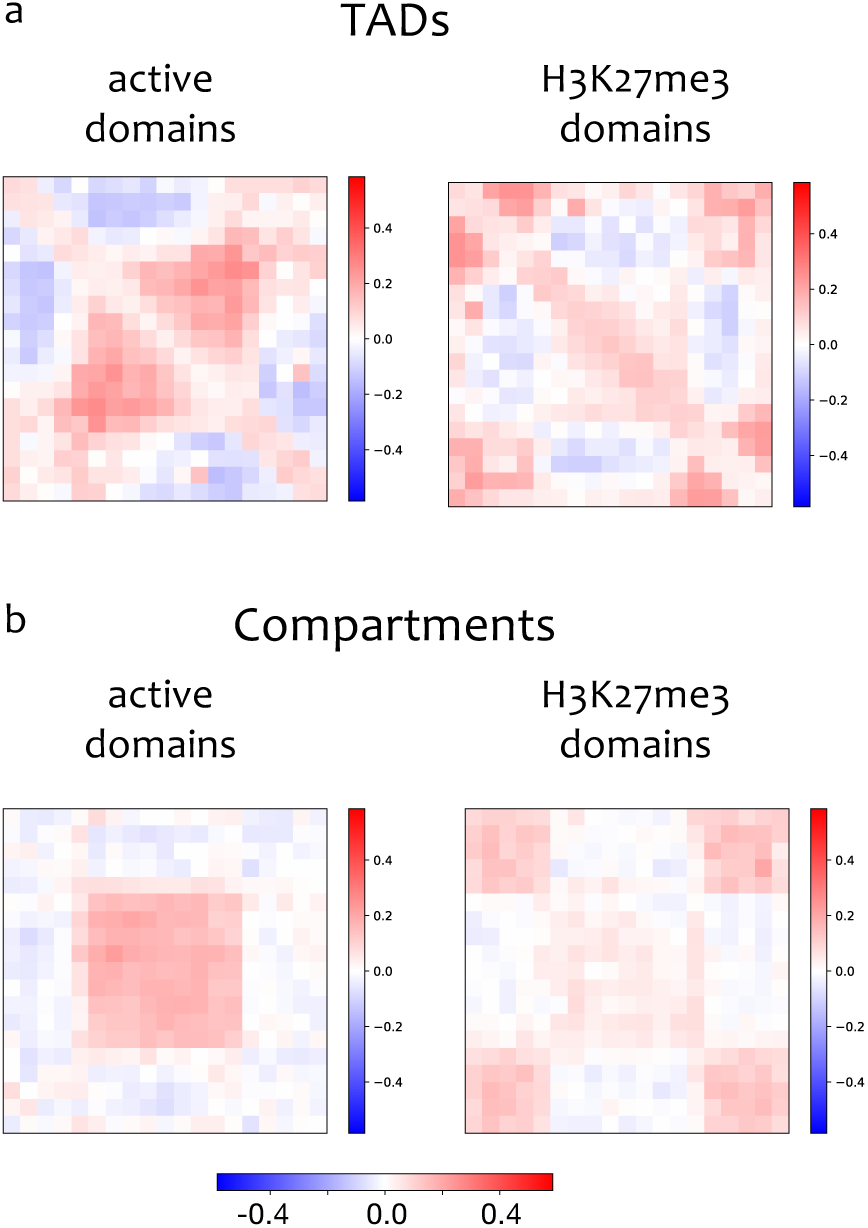
Visualization of TADs and compartments using Hi-C data. Plots are as in Figure 2, but using Hi-C data from ref^7^. (a) TAD enrichment is 1.1 fold for active domains and 1.04 fold for H3K27me3 domains. (b) Compartment enrichment is 1.11-fold for active domains and 1.03 for H3K27me3 domains.

**Supplementary Figure 5.**
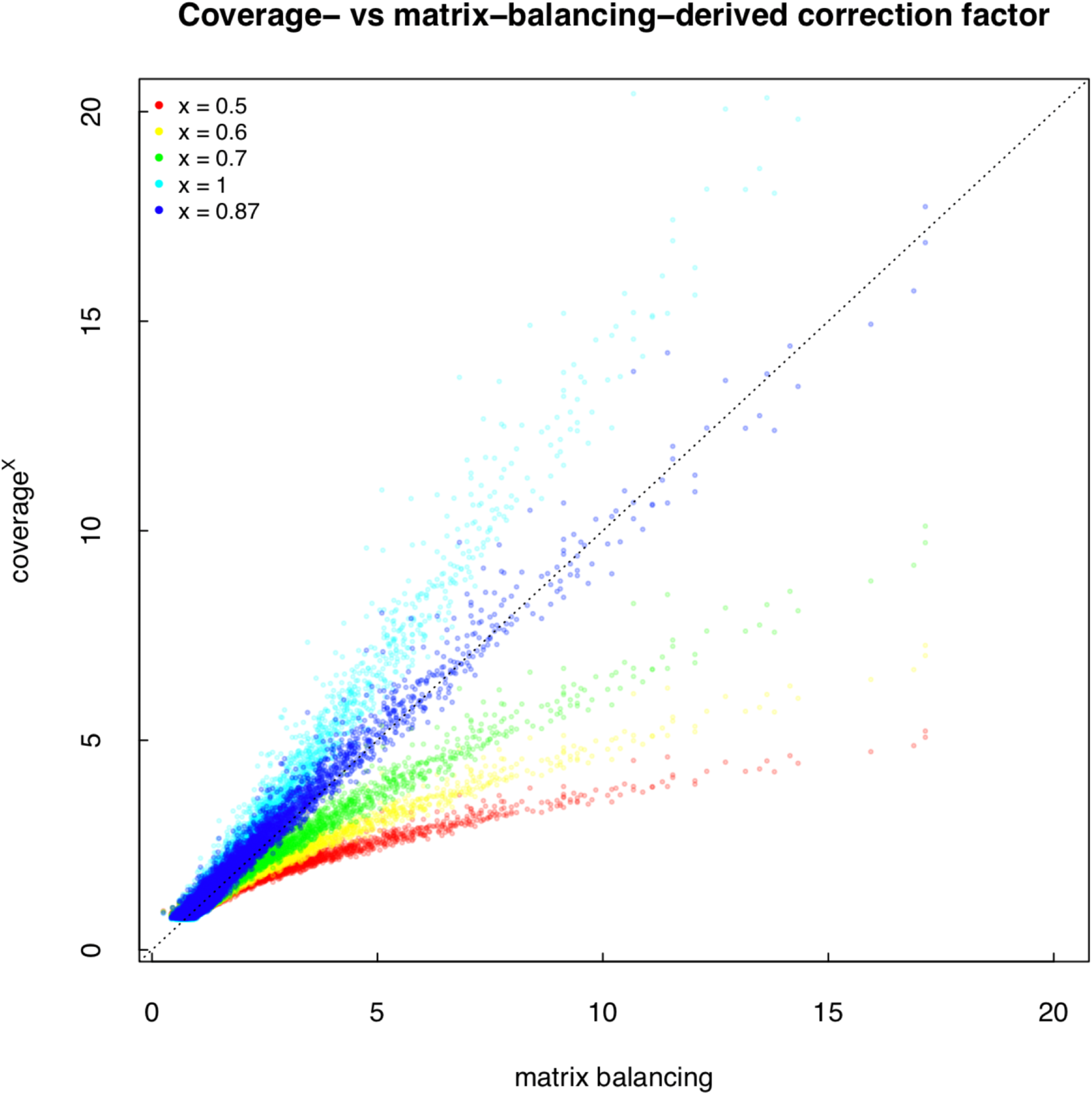
Comparing matrix-balancing-derived correction factors and reciprocal of different exponents of bin coverage In the plot, the x axis gives matrix-balancing derived correction factors (normalised so that the median is 1), whereas y axis gives the reciprocal of different exponents of bin coverage (normalised so that the median is 1).

